## Supplementary File for "Theoretical analysis of cargo transport by catch bonded motors in optical trapping assays"

### 1 Simulation method for single motor

For a cargo transported by a single dynein motor, we consider that the MT is positioned along $y = 0$ and the trap center is set at position $(0, y_o)$. We assume that the cargo starts from the trap center while the motor is attached to the MT at (0,0) at time $t = 0$. We consider $y_o = R + l_o$, where $R$ denotes the radius of the cargo and $l_o$ denotes the restlength of the motor, such that the motor is in a vertical position and the motor is in unloaded mode with an extension equal to $l_o$. This specific choice of initialization also replicates zero restlength scenarios (along the horizontal direction) in our one-dimensional analytical approach. Whenever the motor starts walking it will be subject to a force from the optical trap.

With the force-balance conditions and geometric constraints (Eqs. 7, 8, 9, and 10 in the main text), we perform stochastic simulations of a single dynein-driven cargo transport under optical trap. As time progresses, the motor head attached to the MT walks away from its initial position subject to trap force acting upon it. The force $(f)$ acting on the motor at any time instant is given by $f = k_m(l_m - l_o)$, if $l_m > l_o$ where $l_m = \left[(x_m - x)^2 + y^2\right]^{\frac{1}{2}}$. The motor velocity under load force is assumed to follow the relation, $v_m = v_o(1 - f/f_s)$, where $v_o$ is the single motor velocity at zero load force and $f_s$ is the stall force of the motor. At any simulation time step $\Delta t$, the motor either detaches from the MT with the unbinding rate $\varepsilon$ (Eq. 2 in the main text) or attempts to take a step of size $d$ with probability, $P(\Delta t) = v_m \Delta t/d$. After each motor stepping event the force-balance condition is applied and the values of $x$, $y$, $x_c$, and $y_c$ are obtained from Eqs. 7, 8, 9, and 10. Once the motor unbinds from the MT, the simulation run comes to an end and the corresponding time is considered as the First Passage Time. We averaged all properties over $10^5$ independent simulation runs.

### 2 Simulation method for multiple motors

Transport by multiple motors is simulated only as a 1-D process where the trap center is positioned, by convention, at $x = 0$. Irrespective of the total number of motors present in the system, the simulation always starts with one motor bound. The first motor is bound to the filament with a random choice of the binding site such that the motor binds in a relaxed configuration i.e. force on the motor is zero by $|x_c - x_m| < l_o$ where $x_c$,$x_m$ and $l_o$ are the cargo position, motor position and restlength of the motor respectively. Here, we assume that the motor does not generate any force on compression.

At every time-step the cargo position is recalculated based on the force-balance constraint

$$k_t x_c = \sum_{\text{bound motors m}} \Theta(x_m - x_c - l_o)k_m(|x_m - x_c - l_o|) - \Theta(l_o - x_m + x_c)k_m(x_m - x_c + l_o) \quad (1)$$

where the former term in the sum accounts for forces due to leading motors and the latter term is due to lagging motors. The step function implies that not all bound motors necessarily pull the cargo and that pulling motors, either leading or lagging, need to be determined by the individual motor extensions. Once the cargo position is recalculated, the corresponding motor forces are also recalculated.

As mentioned in the section above, the forces on the individual motors determine the rates by which bound motors stochastically unbind and step forward. However, in the case of multiple motors bound to the cargo, we also have the possibility of binding events i.e. unbound motors can bind with a constant rate $\pi_0$ to the filament.

All motors are updated in parallel at every time-step. Each simulation ends when all motors have unbound and the cargo state is 0. All statistical calculations are done over 20000 cargo trajectories.

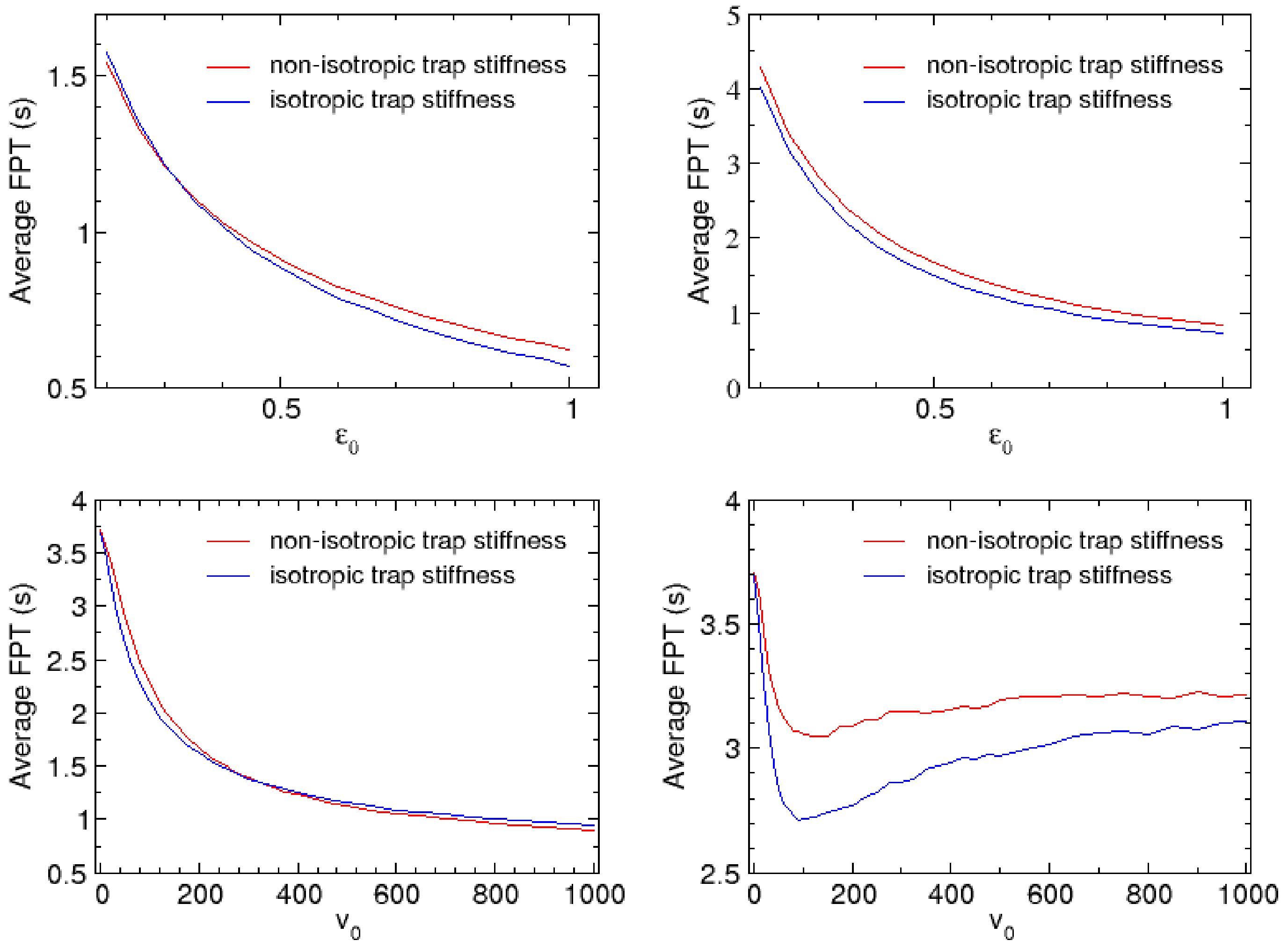


Suppl. Fig. 1: Comparison between the average FPT as a function of unbinding rate (top row) and motor velocity (bottom row) for non-isotropic trap stiffness ( $k_t^y = k_t^x/3 = 0.02/3\ pN/nm = 0.0067\ pN/nm$) with isotropic trap stiffness ( $k_t^y = k_t^x = 0.02\ pN/nm$ ). The left panels denote the slip bond case while the right panels show the catch bond ($\alpha = 18$) scenario. The choice of parameters are the same as Fig. 2 in the main manuscript.

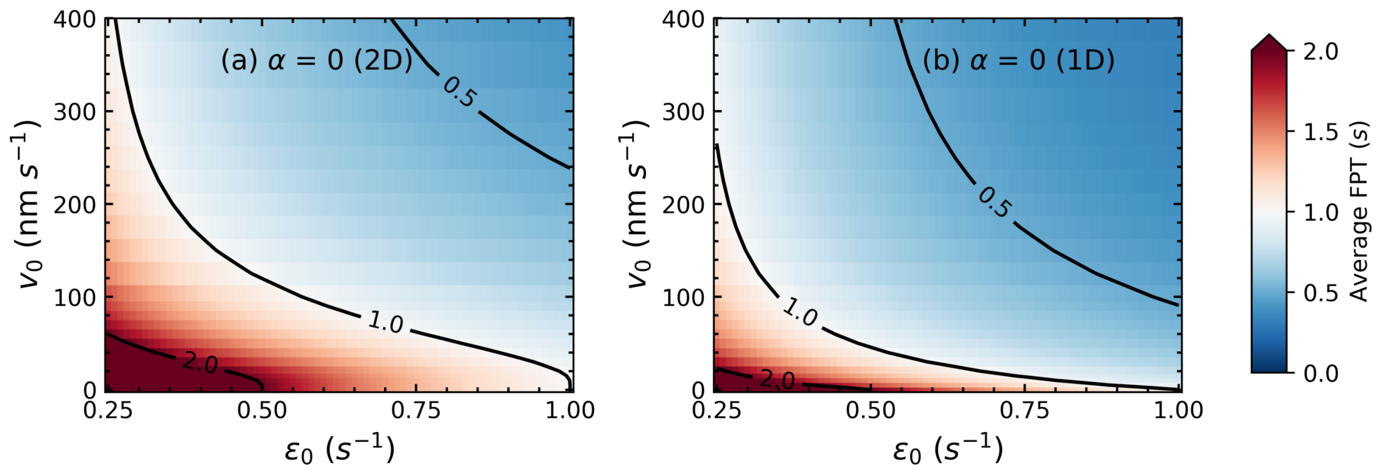


Suppl. Fig. 2: Contour maps for slipbond ($\alpha = 0$) cases in (a) 2D and (b) 1D approach. As is evident from the figure, there is no re-entrant behaviour in case of slipbonds. All other parameter values are the same as in Fig. 2 in the main text.

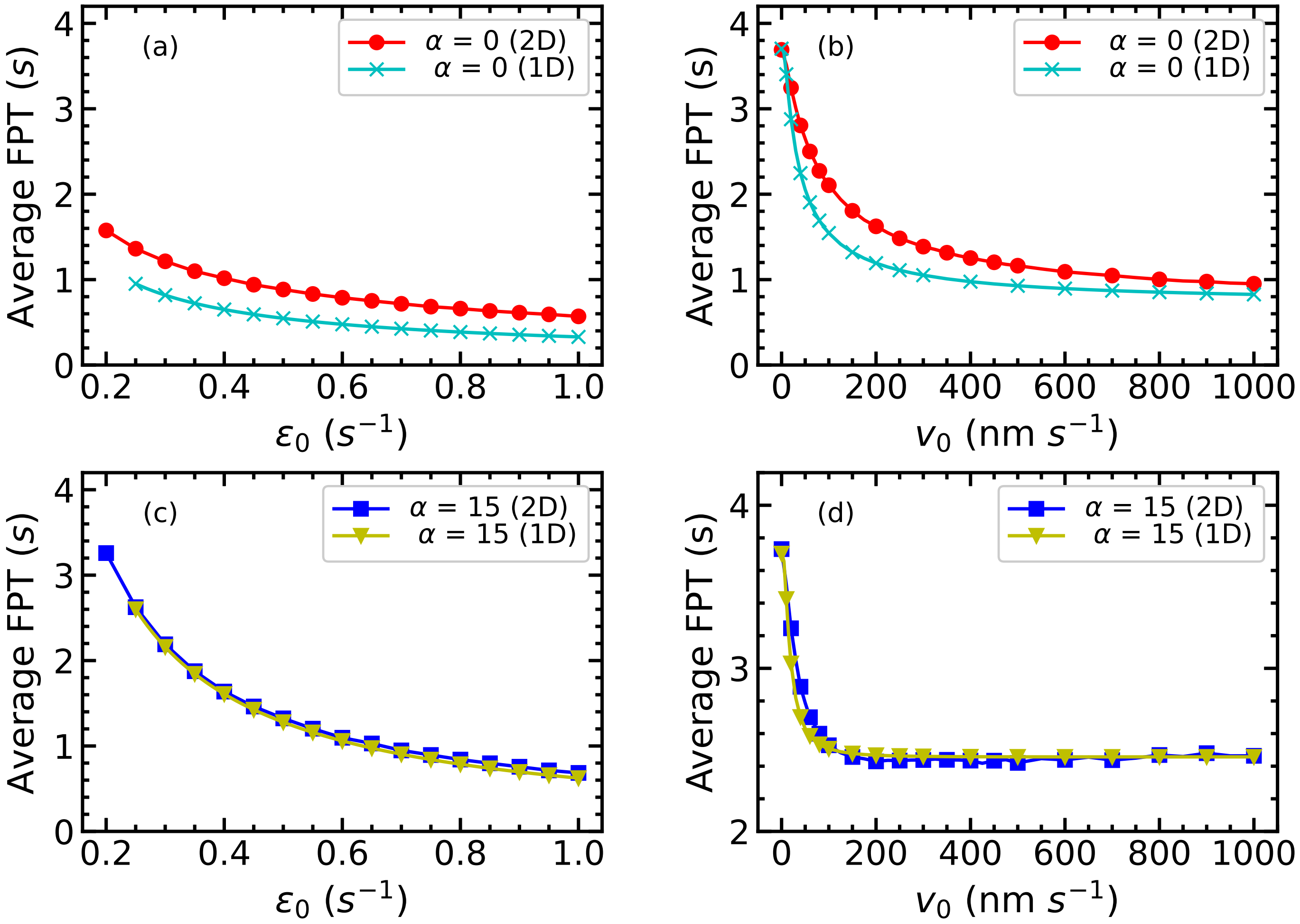


Suppl. Fig. 3: Comparison of average FPT as a function of $\varepsilon$ and $v_0$ obtained in 2D and 1D approaches for slip bond ($\alpha = 0\ k_BT$) and catch bond ($\alpha = 15\ k_BT$). All other parameter values are same as in Fig. 2 in the main text.

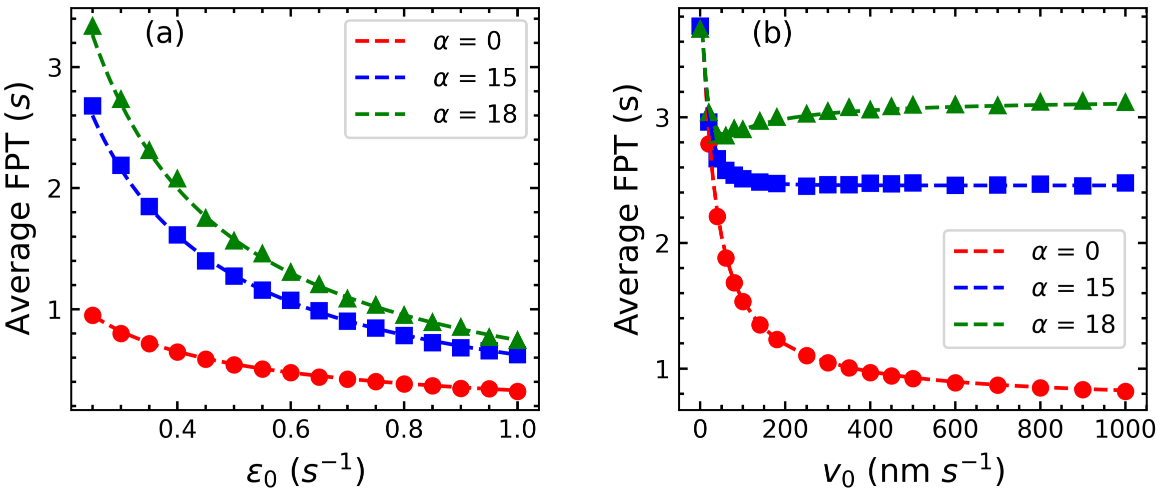


Suppl. Fig. 4: First passage time for single motor cargo transport in 1D. Panels (a) and (b) show the comparison between simulation-generated (dots) and analytically calculated (solid lines) First Passage time $t$ for a single motor. In this case, the motor has a threshold force smaller than its stall force. Red curves correspond to slipbond ($\alpha = 0\ k_BT$) , blue curves to catchbond ($\alpha = 15\ k_BT$) and green curves to Strong catchbond ($\alpha = 18\ k_BT$). All other parameter values are the same as in Fig. 2 in the main text.
